# Mathematical modelling predicts novel mechanisms of stream confinement from Trail/Colec12/Dan in the collective migration of cranial neural crest cells

**DOI:** 10.1101/2025.02.20.639239

**Authors:** Samuel W.S. Johnson, Paul M. Kulesa, Ruth E. Baker, Philip K. Maini

## Abstract

In vertebrate embryogenesis, cranial neural crest cells (CNCCs) migrate along discrete pathways. Analyses in the chick have identified key molecular candidates for the confinement of CNCC migration to stereotypical pathways as Colec12, Trail, and Dan. The effects of these factors on CNCCs *in vitro* are known, but how they confine migration to discrete streams *in vivo* remains poorly-understood. Here, we propose and test several hypothetical mechanisms by which these factors confine cell streams and maintain coherent migration, simulating an expanded agent-based model for collective CNCC migration.

**Results:** Model simulations suggest that Trail enhances adhesion between CNCCs, facilitating movement towards stereotypical migratory pathways, whereas Colec12 confines CNCCs by inducing longer, branched filopodia that facilitate movement down Colec12 gradients and re-connections with streams. Moreover, we find that Trail and Colec12 facilitate the exchange of CNCCs and the formation of CNCC-bridges between adjacent streams that are observed *in vivo* but poorly-understood mechanistically. Finally, we predict that Dan increases the coherence of streams by modulating the speed of CNCCs at the leading edge of collectives to prevent escape.

**Conclusions:** Our work highlights the importance of Trail, Colec12, and Dan in CNCC migration and predicts novel mechanisms for the confinement of CNCCs to stereotypical pathways *in vivo*.

## 1 Introduction

The collective migration of cranial neural crest cells (CNCCs) is a crucial process in vertebrate development that leads to the formation of bone, cartilage, and connective tissues of the face [1, 2]. During embryogenesis, CNCCs emerge from the hindbrain (rhombomere (r) segments r1-7) and migrate dorsolaterally in discrete streams towards the branchial arches (ba1-4), where subsequent cell proliferation and patterning occurs. A crucial aspect of successful migration and subsequent craniofacial patterning is the maintenance of spatial separation between adjacent CNCC streams throughout migration. If mixing between streams occurs, a range of birth defects, termed *neurocristopathies*, may result [3, 4]. As such, a better understanding of the mechanisms governing the confinement of CNCC trajectories to distinct stereotypical migratory pathways could provide pathological insights into craniofacial anomalies and identify therapeutic opportunities to minimise costly repair surgeries [5].

During migration, CNCCs form streams adjacent to rhombomeres r1/2/4/6 that remain separated by regions void of CNCCs adjacent to rhombomeres r3/5 [9] (Figure 1A). In the chick cranial microenvironment, vascular endothelial growth factor (VEGF) is a chemoattractant that facilitates targeted CNCC invasion from r4 to ba2 (Figure 1B). CNCCs express VEGF receptor 2 and Neuropilin 1 receptors and respond to self-induced VEGF gradients (gradients in VEGF generated by CNCC-induced VEGF degradation) [10, 11]. *In vivo* single-cell profiling of chick CNCCs has revealed molecular heterogeneities within migrating streams, with cells at the leading edge of collectives exhibiting upregulated expression of genes that regulate dynamic cell protrusions, ECM degradation, and ECM attachment [12]. Conversely, cells behind the leading edge express cell adhesion genes distinct from leading cells [12]. Such findings suggest that the collective invasion of the branchial arches in the embryonic cranial neural crest is facilitated by a differential response to microenvironmental signals in leading and trailing CNCC populations, wherein leading cells move primarily in response to chemoattractant gradients (such as self-induced gradients in VEGF) and trailing cells move according to cell-cell guidance cues to create loosely-connected streams of CNCCs that populate ba1-4.

**Figure 1:**
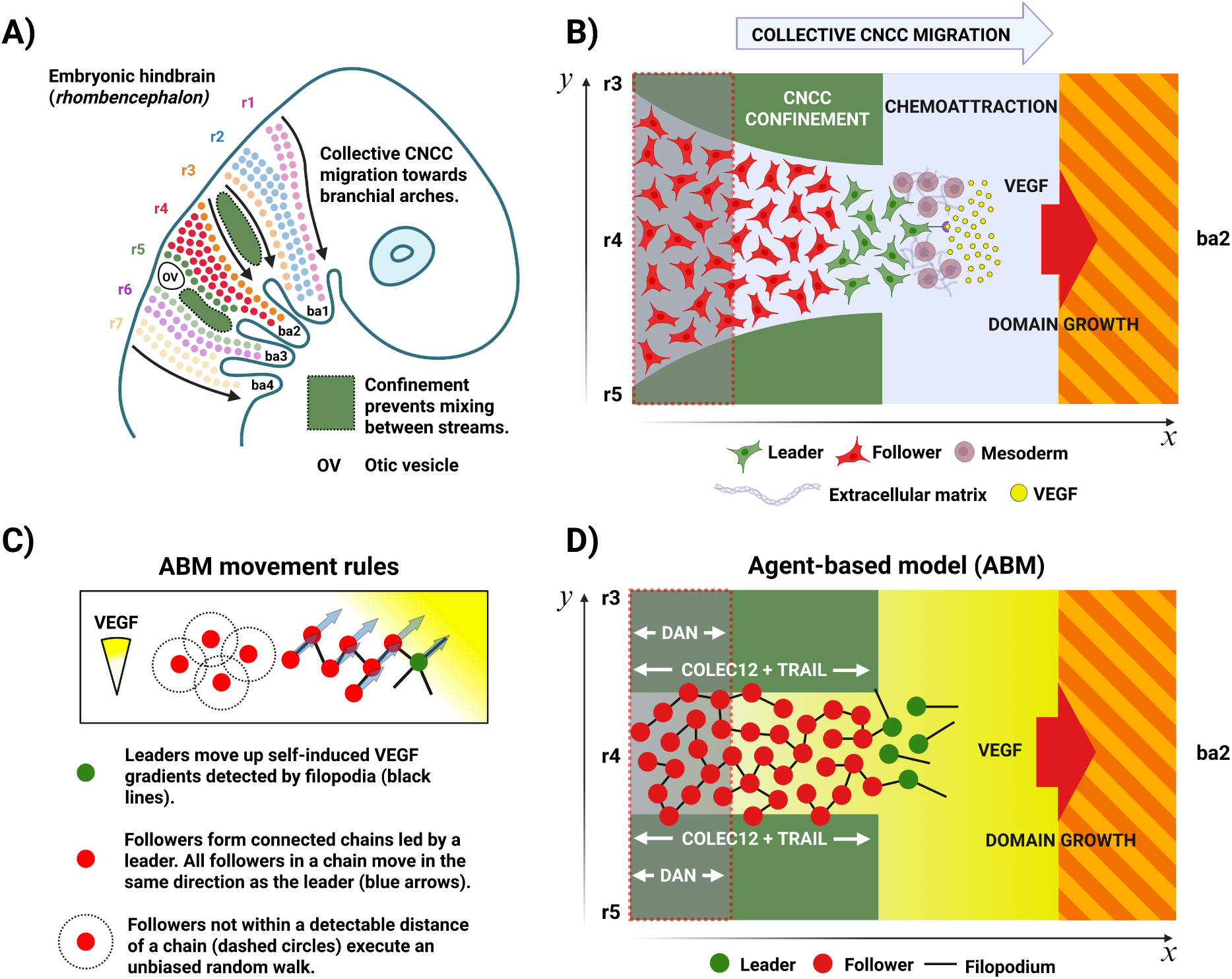
**A**) Schematic of CNCC migration (based on prior works [6, 7, 8]). CNCCs emerge from r1-7 and are sculpted into discrete streams that remain spatially separated during invasion of ba1-4. The majority of CNCCs originate from r2/4/6 and populate ba1/2/3, respectively. CNCCs emerging from r3 and r5 are diverted into the regions adjacent to r2, r4, and r6 due to the presence of inhibitory signals in the paraxial mesoderm adjacent to r3 and r5 (dark green). **B)** Schematic of CNCC migration from r4 to ba2 *in vivo*. CNCCs emerge from r3-r5 and are sculpted into a single stream adjacent to r4 that populates ba2. VEGF guides migration towards ba2, whilst inhibitory signalling in the CNCC-free zones adjacent to r3 and r5 (dark green) prevents migration into the regions adjacent to r3 and r5. Additional inhibitory signalling in the regions proximal to the hindbrain adjacent to r3-5 (translucent grey) modulates CNCC speed after delamination from the rhombomeric hindbrain. **C)** Movement rules in the ABM. Leader cells (green) extend multiple filopodia (black lines) at random orientations to sample local VEGF gradients, and move in the direction of the highest detected positive gradient. Followers (red) form loosely connected chains behind leaders and move in the same direction as the leader at the front of a chain. If a follower moves out of the range of interaction of a chain, it executes an unbiased random walk until a chain is detected. **D)** Schematic of CNCC migration in the model. CNCCs migrate along the r4-ba2 pathway for a period of 24h. Leader cells (green) migrate up self-induced VEGF gradients towards ba2, while follower cells (red) move according to directional cues from cell-cell contacts, facilitating migration as a collective. Trail and Colec12 (dark green) are present for a portion of the regions adjacent to r3 and r5 in the *x*-direction, as observed *in vivo*. Dan (translucent grey) is also present for a portion of the regions adjacent to r3/4/5 and modulates the speed of CNCCs during migration. The domain through which CNCCs migrate grows in the *x*-direction, but remains at a constant width in the *y*-direction.

The factors influencing the exit locations of newly-delaminated CNCCs onto stereotypical migratory pathways in the paraxial mesoderm have been well-studied. Apoptosis is thought to play a role in regulating the number of CNCCs exiting from the odd-numbered rhombomeres, r3 and r5 [13, 14]. However, it is unlikely that apoptosis alone is responsible for maintaining spatial separation between adjacent streams for the duration of migration towards the branchial arches. After emergence from the hindbrain in the regions adjacent to r2/3/4/5, the over-expression of soluble Neuropilin-1/2 Fc in the chick cranial microenvironment has been observed to result in the movement of CNCCs into the CNCC-free zone adjacent to r3 [15]. This observation is suggestive of a role for Neuropilin-1/2 in the early confinement of CNCCs in regions adjacent to the hindbrain (≈ 0 − 200*µ*m from the dorsal midline of the neural tube). However, it is also likely that other factors beyond regions of Semaphorins adjacent to the hindbrain are responsible for confinement in the subsequent stages of migration where CNCCs collectively migrate long distances through the mesoderm between the dorsal neural tube and the branchial arches.

In *Xenopus*, Versican has been shown to confine CNCC collectives by forming inhibitory boundaries in placodal tissues near the branchial arches that prevent the mixing of adjacent streams in late-stage migration [16]. However, Versican loss-of-function experiments show that reduced Versican expression does not result in extensive mixing between adjacent streams *in vivo*, meaning that other factors must also be responsible for the confinement of CNCCs to distinct stereotypical migratory pathways prior to invasion of the branchial arches. Similarly, CNCCs have also been found to avoid regions of EphB2 and Ephrin-B1 expression in the chick cranial microenvironment [17]. However, the expression of EphB2 and Ephrin-B1 lateral to the otic vesicle and ba2 means that these factors are also unlikely to be responsible for the confinement of CNCCs to discrete streams as they migrate away from the hindbrain towards targets in the branchial arches. These observations have led to more recent investigations of the factors expressed in the mesoderm that confine CNCCs to discrete streams for the duration of migration. Expression analyses conducted in the chick cranial microenvironment have highlighted Dan (NBL1, a Dan family BMP antagonist), Trail (tumour necrosis factor-related apoptosis-inducing ligand, CD253), and Colec12 (Collectin-12, a transmembrane scavenger receptor C-type lectin) as factors expressed along, and adjacent to, stereotypical migratory pathways that confine CNCCs to discrete, spatially-separated streams during migration [18, 19]. However, the mechanisms through which these factors confine CNCC movement to stereotypical migratory pathways in the cranial neural crest remain unclear.

Dan has been found to restrict CNCC movement in stripe assays and average cell speed and displacement are reduced *in vitro* when Dan protein is added to the culture media [19]. Furthermore, Dan is expressed in the paraxial mesoderm adjacent to the hindbrain and its concentration is reduced in regions through which CNCCs have previously migrated. These observations are suggestive of a mechanism where migrating CNCCs degrade Dan in the chick, though this hypothesis has not yet been validated experimentally. The addition of Dan into a computational model of CNCC migration showed that cell speed modulation in the presence of Dan is a potential mechanism by which the frequent break-up of collectives observed in prior models can be prevented through a reduction in cell dispersal [19]. However, this model neglected the spatial variations in Dan concentration along the r4-ba2 pathway that are observed *in vivo*, instead assuming that CNCC speed was reduced uniformly across streams exposed to Dan. In reality, it is likely that the reduced concentration of Dan in regions through which CNCCs have previously migrated results in a spatially-varying effect on CNCCs, with a stronger influence on cells at the leading edge where Dan concentration is, on average, higher.

Subsequent analyses in the chick cranial microenvironment have also revealed high expression of Trail and Colec12 in CNCC-free zones adjacent to CNCC streams. *In vitro* assays have highlighted the effects of Trail and Colec12 on chick CNCC morphology, migratory dynamics, and cell-cell interactions [18]. Trail has been found to enhance cell-cell adhesion in chick CNCCs, resulting in clustered amoeboid-like motion rather than movement as individuals, as observed in control media. Moreover, a significant reduction in both the average speed and displacement of CNCCs has been observed in cells cultured in the presence of Trail. Conversely, equivalent assays have highlighted a change in chick CNCC morphology in the presence of Colec12 through the formation of longer, non-retracting filopodial protrusions with increased branching relative to assays in control media. *In vivo*, chick CNCCs in regions of Colec12 expression have also been observed to reverse direction and migrate back towards the neural tube. The mechanism driving this movement back towards the neural tube remains unclear, though it is possible that CNCCs migrate down gradients of Colec12 expression via chemotaxis, which is a well-documented mechanism of migration in CNCC populations [20]. Experiments in the chick show that loss-of-function of either Trail or Colec12 results in the increased deviation of migrating chick CNCCs away from stereotypical migratory pathways and into neighbouring CNCC-free zones. However, as in the case of Dan, the biological mechanisms through which Trail and Colec12 restrict CNCC movement to stereotypical migratory pathways in collective invasion of the branchial arches remain unclear. An ABM for chick CNCC migration has been used to predict that the confinement of CNCCs to the r4-ba2 pathway by factors expressed adjacent to r3/5 is only necessary for the first third of the migratory domain (≈ 400*µ*m) to prevent a breakdown in the collective invasion of ba2 [18].

This prediction guided subsequent experiments, which found the enhanced expression of both Trail and Colec12 for approximately the first 400*µ*m of typical CNCC-free zones adjacent to r3/5. However, this model and other previous models of CNCC migration [12, 19, 21] considered a simplistic representation of inhibitory signalling via the imposition of zero-flux boundary conditions on CNCCs at the axial borders of the regions adjacent to r3/5 and, as such, did not permit an exploration of the mechanisms through which Trail and Colec12 confine CNCC migration to the r4-ba2 pathway *in vivo*.

The discoveries in the chick of factors like Dan [19] and Colec12/Trail [18] that appear to confine CNCC streams and facilitate collective migration are exciting, but lead to further questions regarding the mechanisms by which they achieve confinement. Here, we extend previous computational models of CNCC migration to address the issues that they faced, such as frequent stream-breakage and the necessary, but experimentally unmotivated assumption of zero-flux boundary conditions on cells to confine streams. By using previous experimental observations [18, 19], we construct and test hypothetical mechanisms by which Trail, Colec12, and Dan interact with migrating CNCCs to facilitate collective migration along the r4-ba2 pathway. Using a stochastic ABM for r4-ba2 CNCC migration, we then study the effect of these mechanisms on collective CNCC migration, and hence, aim to provide new insights into the complex interplay between, and relative importance of, Trail, Colec12, and Dan in maintaining collective migration along stereotypical pathways in the cranial neural crest. The model couples an existing stochastic agent-based model (ABM) for chick CNCC r4-ba2 migration [12] to a continuum partial differential equation (PDE) model for the dynamics of factors expressed adjacent to and along stereotypical migratory pathways, and builds on prior iterations of the model by explicitly integrating the effects of chemical signalling on cell speed, direction, and cell-cell interactions.

## 2 Results

The ABM for CNCC migration focuses on the migration of CNCCs through a region adjacent to r4, the r4-ba2 pathway, that is enclosed by CNCC-free zones adjacent to r3 and r5 (Figure 1B). The model is a two-dimensional representation of *in vivo* migration through the three-dimensional mesenchyme of the chick embryo, and as such, may be taken to represent the migration of a single layer of CNCCs within a stream. Cells are represented as discrete two-dimensional non-overlapping discs whose migration depends on the concentration of VEGF, Dan, Trail, and Colec12 in their neighbourhood. *In vivo*, CNCCs migrate along curved trajectories [22]. Though the effect of domain curvature on migration is unknown, for this ABM, it has recently been shown that for a purely deterministic model, curvature does not significantly impact cell motion [23]. As such, the r4-ba2 pathway is approximated as a two-dimensional rectangle in which VEGF is present everywhere and Dan is present for a certain proportion of the pathway in the *x*-direction (Figure 1D). Hence, the r4-ba2 pathway is assumed to grow in length in a spatially uniform manner from approximately 300*µ*m to 1100*µ*m over a period of 24h and its width is a constant value of 120*µ*m throughout migration, corresponding to the anterior-to-posterior length of r4 in the chick. The assumption of spatially uniform domain growth is made for computational simplicity, though it has been observed that mesoderm grows non-uniformly in both space and time during chick head morphogenesis [22]. Relaxing this model assumption is a subject of future work. The r4-ba2 pathway is also enclosed by two CNCC-free zones adjacent to r3 and r5, each of width 120*µ*m, in which VEGF is expressed everywhere and Trail, Colec12, and Dan are present for a certain proportion of the domain in the *x*-direction, and that grow in length at the same rate as the r4-ba2 pathway. For the chick embryo, there is currently no explicit discussion of VEGF expression adjacent to r3/5 in the experimental literature. Here, we assume that VEGF is expressed in these regions as this is likely to drive the invasion of leader cells into CNCC-free zones and hence, presents the best test of robustness for our proposed mechanisms of CNCC confinement. As in prior ABMs of CNCC migration [12, 18, 19], cells adopt one of two phenotypes – leader or follower. In leader-follower models of CNCC migration, cells at the leading edge of collectives move up self-induced VEGF gradients sensed by filopodial protrusions. Cells behind the leading edge, where VEGF gradients are not detectable due to the degradation and dilution of VEGF by CNCCs and tissue growth, respectively, instead form chains of cells close to each other, in which all follower cells receive the same directional cues from a leader cell at the front of a chain (Figure 1C). In the ABM, a follower cell joins a chain upon the detection of another cell currently in a chain. Once in a chain, follower cells move in the same direction as the leader cell at the front of their respective chain, to create a stream of cells that migrate as a collective. In the chick cranial neural crest, VEGF signalling induces the expression of genes associated with a leader cell phenotype [21]. To reflect this in the ABM, we implement a mechanism of phenotype switching, wherein a follower cell that overtakes a leader cell adopts a leader phenotype and the leader cell that is overtaken adopts a follower phenotype, thus ensuring that leader cells remain at the leading edge of collectives where VEGF concentration is highest, on average. This mechanism is a computationally simple abstraction of VEGF-induced phenotype switching that has been used in a prior model of CNCC migration [12] and in many models of CNCC migration produces qualitatively similar migratory profiles to those in which phenotype switching depends on time-averaged VEGF exposure [21]. This mechanism also fixes the number of leader cells in a simulation, which is another common simplification in models of CNCC migration. A full description of the ABM and parameter values can be found in Section 4 and videos of example simulations can be found in Supplementary Information S1/S2/S3.

### 2.1 Trail and Colec12 confine CNCCs to the r4-ba2 pathway in collective migration

Expression profile analyses of the paraxial mesoderm through which CNCCs migrate have revealed enhanced levels of Trail and Colec12 expression in typically CNCC-free zones adjacent to r3 and r5. Furthermore, *in vitro*, CNCCs have been found to avoid regions of either Trail or Colec12 expression [18]. These findings suggest that both Trail and Colec12 play a role in the confinement of CNCCs to stereotypical migratory pathways during migration.

*In vitro* experiments have shown that CNCCs exposed to Colec12 protein extend longer, more branched protrusions that retract at a slower rate than in control media [18]. To investigate the effects of Colec12 on the spatial confinement of CNCCs, we used the ABM to study CNCC migration along the r4-ba2 pathway with Colec12 present in the CNCC-free zones adjacent to r3 and r5. In the ABM, Colec12 is present for a certain proportion of the CNCC-free zones adjacent to r3 and r5 (Figure 2A). The proportion of CNCC-free zones for which Colec12 is present remains constant during domain growth, such that the distance over which Colec12 is present in CNCC-free zones increases as the domain is extended. CNCCs that encounter Colec12 extend filopodial protrusions that are 1.5 times longer than at baseline. We choose to represent the increased branching of filopodia in CNCCs exposed to Colec12 by the extension of five times the number of filopodia in a given time interval. Furthermore, we represent the slower retraction of filopodia by CNCCs by increasing the time interval over which cells extend new filopodia by a factor of 10 (such that a given filopodium remains extended for a longer period of time). A visual summary of the effects of Colec12 protein on CNCCs in the ABM can be seen in Figure 2C and a full description of the parameterisation and representation of the effects of Colec12 in the ABM can be found in Section 4.

**Figure 2:**
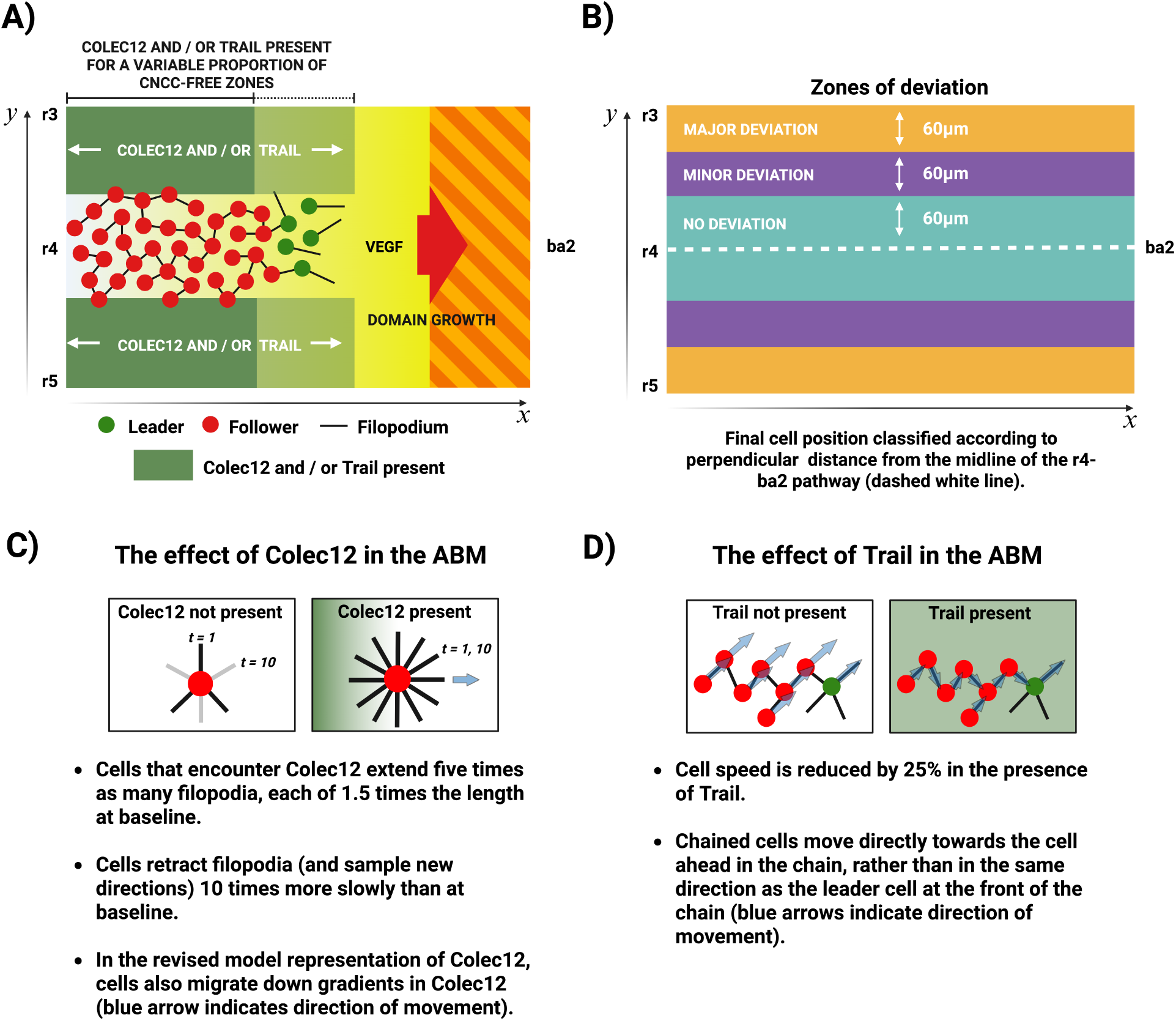
**A**) Schematic of the expression patterns of Trail and Colec12 in Section 2.1. Trail and / or Colec12 are expressed for a certain proportion of the CNCC-free zones adjacent to r3/5 in the *x*-direction, which is varied to study the effects of Trail and Colec12 expression on CNCC confinement. In the initial computational experiments where Trail and Colec12 are expressed for variable proportions of CNCC-free zones, Dan is not expressed anywhere in the domain and VEGF is expressed everywhere in the domain. **B)** Schematic showing zones of deviation in the ABM. The final position of CNCCs in the domain is classified by their perpendicular distance from the midline of the r4-ba2 pathway after 24h of migration. If a cell remains in the r4-ba2 pathway after 24h, it lies in a zone of no deviation. Cells in CNCC-free regions within 60 − 120*µ*m of the mid-line of the r4-ba2 pathway after migration are classified as having exhibited a minor deviation. All other cells are classified as having exhibited a major deviation in migration. **C)** Schematic for the effects of Colec12 in the ABM. CNCCs that encounter Colec12 extend five times the number filopodial protrusions, that are each 1.5 times longer in length than at baseline. Filopodia are also retracted 10 times more slowly than at baseline. In the revised model representation of the interaction between Colec12 protein and CNCCs, CNCCs also undergo chemotaxis down concentration gradients in Colec12, driving movement away from regions of Colec12 expression back towards the r4-ba2 pathway. **D)** Schematic for the effects of Trail in the ABM. Movement speed is reduced by 25% when cells encounter Trail. Follower CNCCs exposed to Trail also move directly towards the cell in front of them in a chain to represent an increase in cell-cell adhesion.

In order to quantify the extent to which Colec12 confines CNCCs to the r4-ba2 pathway in model simulations, we define three regions of the r4-ba2 pathway and adjacent CNCC-free zones according to their distance from the central mid-line of the r4-ba2 pathway (Figure 2B). After 24h of migration, we classify CNCC positions in zones of no, minor, or major deviation according to the end point of their trajectory in the domain.

Simulations of the ABM show that CNCC interactions with Colec12 sub-regions have a minimal effect on the overall confinement of migrating CNCCs to the r4-ba2 pathway. By varying the proportion of CNCC-free zones for which Colec12 is present (Figure 2A), we found no correlation between the proportion of CNCC-free zones for which Colec12 is present and the extent to which CNCCs remain confined to the r4-ba2 pathway for the 24h period of migration (Figure 6). For all distances of Colec12 expression in CNCC-free zones, we found that approximately 35% of CNCCs lay in regions of minor or major deviation after migration ended. Furthermore, we found that this lack of confinement to the r4-ba2 pathway led to the frequent break-up of collectives and a subsequent breakdown in collective migration. This finding is suggestive of an additional mechanism of interaction between Colec12 and CNCCs that is responsible for their confinement during migration.

Motivated by the observation that CNCCs exposed to Colec12 *in vivo* frequently change direction and migrate back towards the neural tube [18], we added a mechanism of chemotaxis in CNCCs exposed to Colec12 protein, such that they migrate down concentration gradients of Colec12 in the model (Figure 2C). When this mechanism of interaction was added to the model representation of Colec12, a strong dependence between the extent of CNCC confinement to the r4-ba2 pathway and the proportion of CNCC-free zones for which Colec12 is expressed in CNCC-free regions emerged. We found that as the proportion of CNCC-free zones for which Colec12 is expressed increases, the overall confinement of CNCCs to the r4-ba2 pathway after the 24h period of migration increases from approximately 65% confinement for no Colec12 expression in CNCC-free zones to approximately 95% confinement when Colec12 is expressed for the full length of CNCC-free zones (Figure 3A). Based on these observations, we hypothesise that chemotaxis down gradients in Colec12 protein is the primary mechanism through which Colec12 confines CNCCs to the r4-ba2 pathway during migration. The morphological effects of Colec12 on CNCCs observed *in vitro* may, therefore, be interpreted as mechanisms that increase the likelihood of gradient detection through the extension of longer, more branched filopodia.

**Figure 3:**
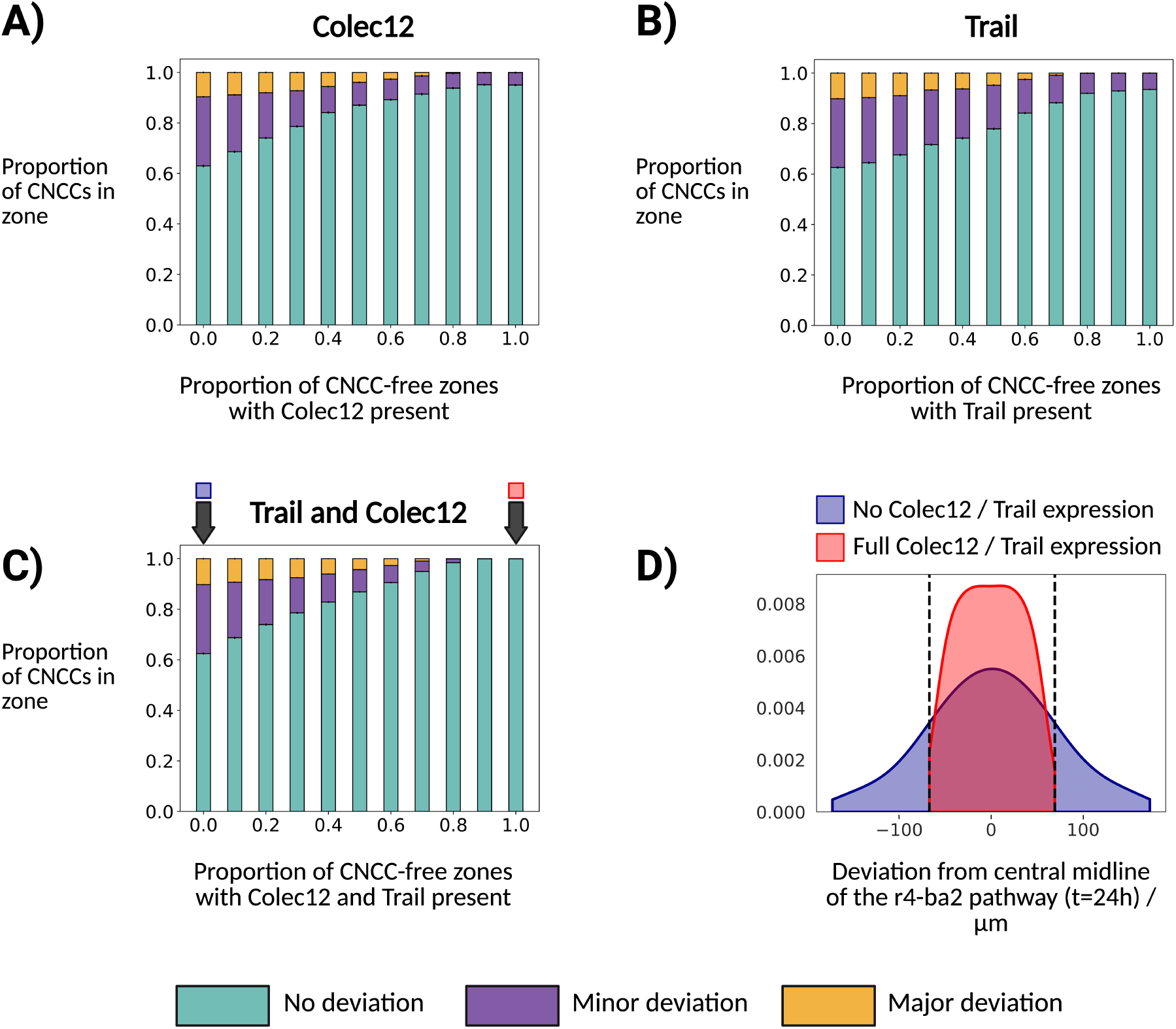
**A)-C**) The confinement of migrating CNCCs to the r4-ba2 pathway as a function of the fraction of CNCC-free zones for which Colec12, Trail, or Colec12/Trail are present. In all simulations, Dan is not present anywhere in the domain and VEGF is present everywhere in the domain. **D)** Truncated kernel density estimates for the distribution of final CNCC *y* coordinates relative to the mid-line of the r4-ba2 pathway when Trail and Colec12 are not expressed and are expressed for the full length of CNCC-free zones. Dashed lines indicate the maximum and minimum deviations when Trail and Colec12 are expressed for the full length of CNCC-free zones. Both kernel density estimates are displayed only between the maximum and minimum values in each respective data set to avoid displaying non-zero densities for distances beyond extremal data points. The results in each plot are averaged over 500 simulations.

To study the effect of Trail expression in CNCC-free zones adjacent to r3/5, we conducted a similar investigation by varying the fraction of CNCC-free zones for which Trail is present. *In vitro*, Trail reduces the speed at which CNCCs move and increases adhesion between CNCCs, resulting in an amoeboid-like movement [18]. To represent the effects of Trail on CNCCs in the ABM, we reduce cell speed in the presence of Trail protein by 25% and alter the mechanism through which follower CNCCs move when they encounter Trail. In the absence of Trail, follower cells in the ABM move by forming chains of cells with a leader at the front (Figure 1C). If a follower cell lies in a chain at the time of movement, it moves in the same direction as the leader cell at the front of its respective chain. In regions of Trail expression, we represent an increase in cell-cell adhesion by altering the mechanism of follower cell movement, such that follower cells instead move directly towards the cell in front of them in their respective chain, directing CNCC movement towards areas of higher cell density, on average. A visual summary of the effects of Trail expression on CNCCs in the ABM can be seen in Figure 2D and a full description of the parameterisation and representation of Trail expression in the ABM can be found in Section 4.

As in the case of our modified model for Colec12 expression in CNCC-free zones, we find a strong dependence between the proportion of CNCCs confined to the r4-ba2 pathway at 24h and the fraction of CNCC-free zones adjacent to r3 and r5 for which Trail is expressed. Model simulations suggest that increasing the length over which Trail is expressed in CNCC-free zones increases the overall confinement of CNCCs to the r4-ba2 pathway from approximately 65% confinement for no Trail expression to approximately 95% confinement for expression along the entire length of CNCC-free zones (Figure 3B). These observations suggest that the effects of Trail on CNCCs noted *in vitro* are mechanisms through which Trail may confine CNCCs to the r4-ba2 pathway during migration. By increasing cell-cell adhesion, Trail may drive CNCC movement into regions of high cell density, and hence, towards the stream along the r4-ba2 pathway.

We concluded our investigation of the effects of Trail and Colec12 expression in CNCC-free zones by investigating the combined effects of both Trail and Colec12 expression in CNCC-free zones adjacent to r3 and r5 on the confinement of CNCCs to the r4-ba2 pathway. As in our prior investigations of the effects of either Trail or Colec12 expression in CNCC-free zones, we conducted simulations in which both Trail and Colec12 are expressed for a fraction of the CNCC-free zones adjacent to r3/5. Simulations of this model reveal that the proportion of CNCCs confined to the r4-ba2 pathway with both Trail and Colec12 expressed is higher than in the cases of either Trail or Colec12 expression in CNCC-free zones (Figure 3C). We find that the confinement of CNCCs to the r4-ba2 pathway increases from 65% for no Trail or Colec12 expression in CNCC-free zones adjacent to r3 and r5 to almost 100% confinement for the expression of both Trail and Colec12 along the full length of CNCC-free zones. The expression of both Trail and Colec12 in the CNCC-free zones adjacent to the r4-ba2 pathway is, therefore, a mechanism that mitigates against large migratory defects by preventing the invasion of CNCC-free zones (Figure 3D).

To summarise, model simulations of Trail and/or Colec12 expression in CNCC-free zones adjacent to stereotypical migratory pathways suggest that Trail and Colec12 are complementary factors for the confinement of CNCCs to stereotypical migratory pathways during migration. In simulations, the confinement of CNCCs to stereotypical migratory pathways increases with the fraction of CNCC-free zones for which either factor is expressed. Furthermore, the confinement of CNCCs to stereotypical migratory pathways in model simulations is maximised when both Trail and Colec12 are present in CNCC-free zones.

### 2.2 CNCC-induced degradation of Dan increases stream coherence through differential CNCC speed modulation

Dan is a bone morphogenetic protein (BMP) antagonist that has been found to exhibit a dynamic expression profile along stereotypical migratory pathways in the chick cranial neural crest [19]. Prior to migration, Dan is present throughout the regions parallel to the body axis adjacent to r1-7. However, after CNCCs populate the branchial arches, Dan concentration is lower in regions through which CNCCs have previously migrated. This observation suggests that Dan may be degraded by CNCCs during their migration, though this has not yet been verified experimentally. *In vitro*, CNCCs avoid regions of Dan expression and move more slowly when they encounter Dan protein, suggesting that Dan may restrict the movement of CNCCs during migration by lowering their average speed and displacement. Simulations of a previous computational model support this hypothesis by suggesting that Dan prevents the escape of cells at the leading edge of CNCC collectives by modulating their speed [19], which is necessary to ensure proper development in the chick [12]. However, this computational model did not fully account for the dynamic concentration profile of Dan, in that the potential degradation of Dan by CNCCs was not accounted for and its concentration was assumed to be spatially uniform in a subregion of the domain through which migration occurs. This in turn, did not permit a study of the effects of preferential, rather than uniform, speed moderation in cells at the leading edge of collectives which may arise due to interactions between Dan and CNCCs.

In order to investigate the mechanisms through which Dan promotes collective migration in the cranial neural crest, we represent Dan as a degradable, non-diffusive chemical that is initially present for approximately the first one-third of the regions adjacent to r3/4/5 (Figure 1D). As CNCCs move through the domain, they locally degrade Dan to produce a dynamic concentration profile in which the concentration of Dan is lower in regions through which CNCCs have previously migrated (Figure 4A). To represent the modulation of CNCC speed by Dan observed *in vitro* [19], we reduce the speed of CNCCs that encounter Dan by a factor proportional to the concentration of Dan at a cell’s current position (Section 4.4). We then repeated previous model simulations in which Trail and Colec12 are expressed for a certain fraction of CNCC-free zones adjacent to the r4-ba2 pathway (Figure 3A) to study the effect of Dan on the frequency of leader escape during collective migration.

**Figure 4:**
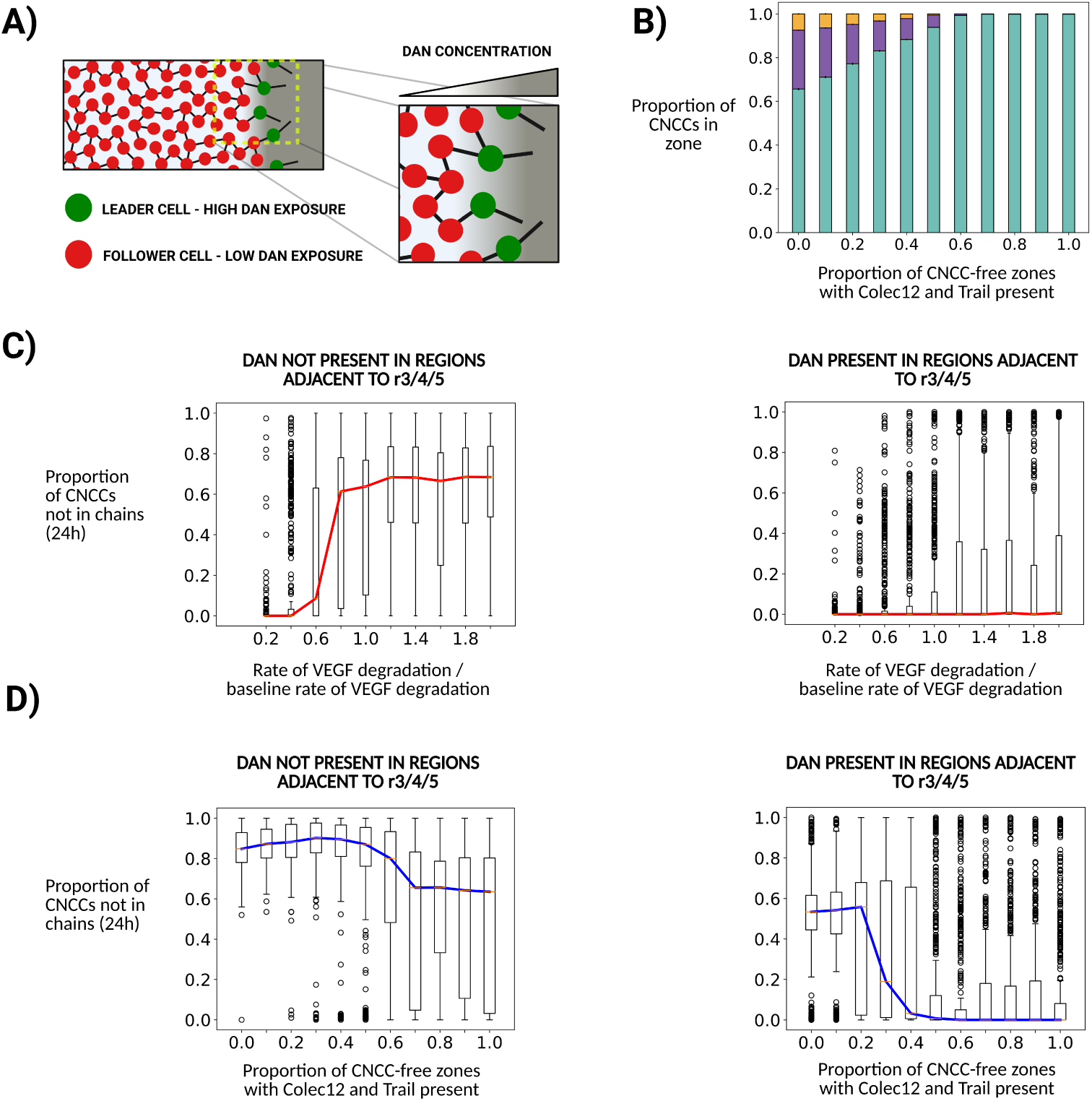
**A**) Schematic of leader cell speed modulation by Dan in CNCC migration. CNCCs at the leading edge of collectives invade regions in which Dan protein concentration is high, leading to a reduction in their average speed and displacement. Trailing cells in collectives migrate through regions of lower Dan concentration and hence move with a higher average speed, thus preventing a loss of contact with cells closer to the leading edge of streams. **B)** The confinement of migrating CNCCs to the r4-ba2 pathway as a function of the fraction of CNCC-free zones for which Trail and Colec12 are present with Dan present for approximately the first one-third of the r4-ba2 pathway and VEGF present everywhere in the domain. **C)** Boxplots of the proportion of CNCCs in chains after 24h of migration as a function of VEGF degradation rate. When Dan is not present, high rates of VEGF degradation (and hence, the formation of VEGF gradients along which leader cells migrate) lead to a break-up of streams due to leader escape. When Dan is present in the first one-third of the r4-ba2 pathway, migration is robust to rapidly induced VEGF gradients. Here, Trail and Colec12 were expressed for the full length of the CNCC-free zones adjacent to r3/5. Red lines indicate the median proportion CNCCs in streams after 24h, boxes indicate the upper and lower quartile values, and circles indicate outliers. The baseline rate of VEGF degradation in simulations is 2.75 × 10^2^*µ*m^2^/h. **D)** Boxplots of the proportion of CNCCs in chains after 24h of migration as a function of the proportion of CNCC-free zones for which Trail and Colec12 are expressed. When Dan is not present in the domain, the median fraction of CNCCs not in streams after migration ends at *t* = 24h remains above a value of 0.6 for all Trail/Colec12 expression lengths along CNCC-free zones adjacent to the r4-ba2 pathway. For Dan expression along the first one-third of the r4-ba2 pathway, the proportion of CNCCs in streams after migration ends at *t* = 24h exhibits phase transition-like behaviour as the length of CNCC-free zones for which Trail/Colec12 are expressed increases. When Trail/Colec12 are expressed for at least half of CNCC-free zones in the proximal-to-distal direction, all CNCCs are in chains when migration ends, on average. Blue lines indicate the median proportion of CNCCs in streams after 24h, boxes indicate the upper and lower quartile values, and circles indicate outliers. The results in figures **B)**, **C)**, and **D)**, are each averaged over 500 simulations.

Model simulations show that the presence of Dan in the regions through which CNCCs migrate is essential for collective migration to occur by preventing the break-up of collectives. Furthermore, we find that the degradation of Dan is a mechanism that confers robustness on migration, in that dynamic Dan concentration profiles increase the range of CNCC-induced VEGF degradation rates for which collective migration is preserved (Figure 4C). When Dan is degraded by CNCCs, we find that it functions as a preferential modulator of speeds at the leading edge of CNCC collectives (Figure 4A), such that it prevents the break-up of collectives when leader cell movement along VEGF gradients occurs too rapidly for coherent stream migration to occur in follower populations. In simulations where Dan is not present in the regions adjacent to r3/4/5, at least 60% of CNCCs are not in chains when migration ends after a period of 24h, on average (Figure 4D). However, when Dan is present for the first one-third of the r4-ba2 pathway we find that for Trail/Colec12 expression distances of greater than approximately one-half of the CNCC-free zones adjacent to r3/5, all CNCCs remain in streams throughout migration, on average. The addition of Dan expression into the model also reproduces the observation from a previous computational model that collective migration towards the branchial arches can occur for only partial expression of Trail and Colec12 in the regions adjacent to r3/5 [18] (Figure 4B).

In summary, model simulations suggest that Dan expression along the r4-ba2 pathway is necessary to prevent the break-up of CNCC streams during migration due to the escape of cells at the leading edge of collectives. As CNCC collectives move along stereotypical migratory pathways, cells at the leading edge of collectives invade regions of high Dan expression, such that their speed is significantly reduced. Due to the degradation of Dan by CNCCs in the ABM, trailing cells migrate into regions of lower Dan concentration, such that their speed of movement is higher than leading cells, on average, thus preventing the escape of leader cells that results in a breakdown of collective migration. When Dan, Trail, and Colec12 are present in their typical patterns of expression (Figure 1C), CNCC migration in the model is both highly confined to the r4-ba2 pathway, and coherent in the sense that all cells, on average, remain in streams throughout migration towards ba2.

### 2.3 Trail and Colec12 expression in CNCC-free zones facilitate the exchange of CNCCs between adjacent collectives

Timelapse imaging of chick cranial neural crest migration has revealed the exchange of a small number of CNCCs between streams emerging from regions adjacent to r4 and r6 [24]. Whilst streams adjacent to r4 and r6 remain largely separated during invasion of ba2 and ba3 *in vivo*, a small number of CNCCs within these streams invade the region lateral to the otic vesicle (OV) lying between the typical migratory pathways from r4 and r6 to ba2 and ba3, respectively. This results in the exchange of a small number of CNCCs between the streams along the r4-ba2 and r6-ba3 pathways over distances of 50 − 100*µ*m, without the breakdown of collective migration. Furthermore, npn2 or sema3F loss-of-function assays in the chick have demonstrated the formation of single-cell wide bridges between streams invading ba1 and ba2, without large-scale disruption to migration [25]. Hitherto, the mechanisms underpinning these phenomena have not been explained through experiments or computational modelling.

Using the ABM for CNCC migration, we investigated the effects of Trail and Colec12 expression in the CNCC-free zone adjacent to r3 on the exchange of cells between streams invading ba1 and ba2. In order to study the exchange of CNCCs between streams, we modified the model geometry to include two streams adjacent to r2 and r4 that are separated by a CNCC-free zone adjacent to r3 in which Trail, Colec12, and Dan are expressed (Figure 5A). As before, VEGF is present everywhere in the domain [10] and Dan is expressed for approximately the first one-third of the regions adjacent to all rhombomeres. In this revised model geometry, Trail and Colec12 are also expressed for the full length of the regions adjacent to r1/5 along the migratory pathway. The expression of Trail and Colec12 in the regions adjacent to r1/5 has not been confirmed experimentally, but is included to keep streams confined to the regions adjacent to r2/4 during migration. However, the mechanistic inferences made in this section depend only on the expression of Trail/Colec12 in the region adjacent to r3, which has been observed *in vivo* [18].

**Figure 5:**
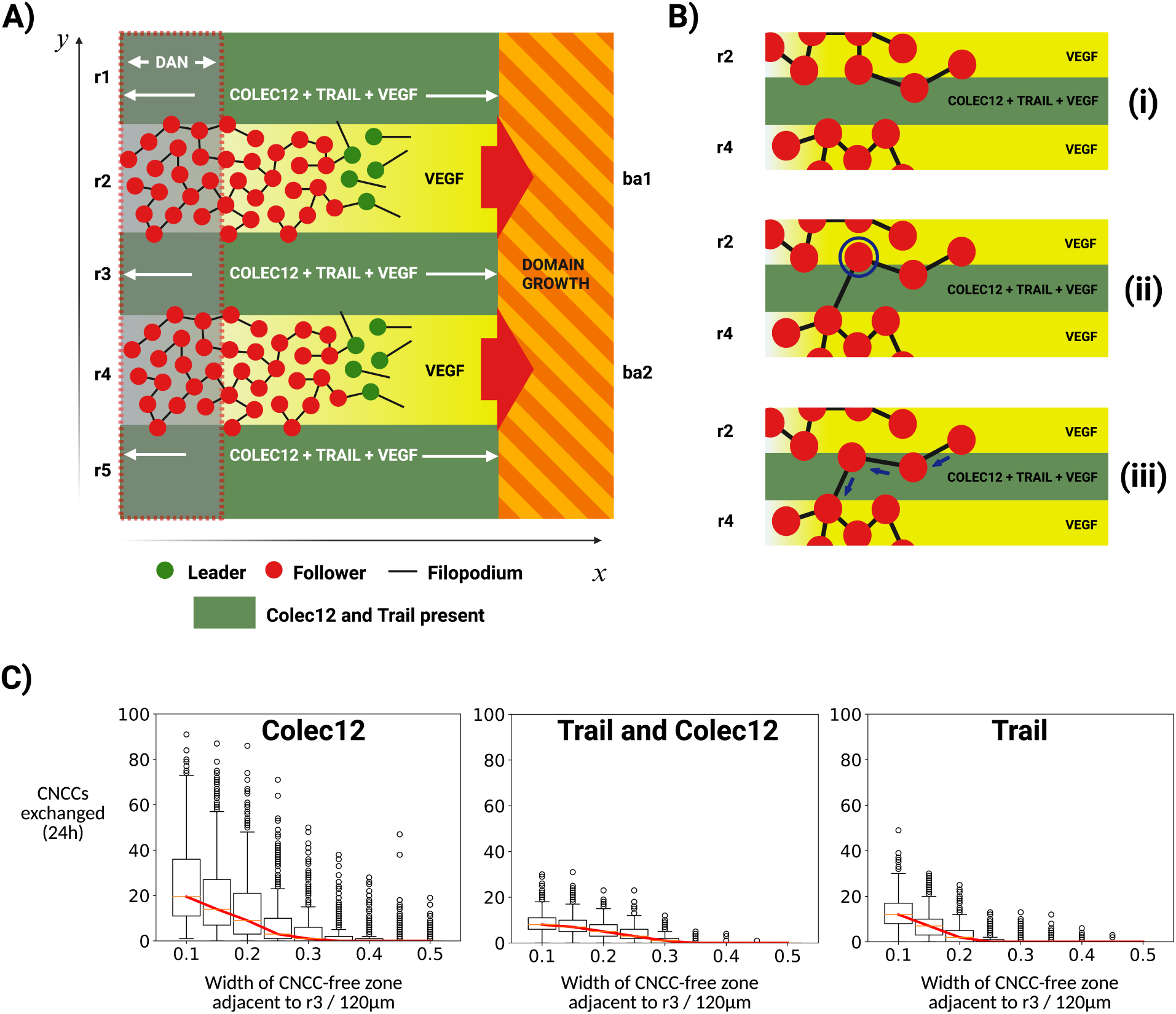
**A**) Revised model geometry for studying cell exchange between adjacent streams. **B)** Mechanism for the formation of CNCC-bridges between adjacent collectives for when Trail (or both Trail and Colec12) is expressed in CNCC-free zones. **(i)** Cells migrate in distinct streams towards ba1/2. **(ii)** A cell (circled) loses contact with cells in its current stream and detects a cell in the adjacent stream with filopodial protrusions. **(iii)** The presence of Trail in the CNCC-free zone separating streams adjacent to r2 and r4 facilitates cell-cell adhesion that drives the movement of the cell across the mid-line of the CNCC-free zone adjacent to r3 (dashed-black). Other cells in the presence of Trail also begin to move towards one another (blue arrows), forming a chain of cells that invade the region adjacent to r3. **C)** Boxplots of CNCC exchange frequency in the model as a function of the width of the region adjacent to r3 for Trail, Trail/Colec12, and Colec12 expression in the CNCC-free zone adjacent to r3. Red lines indicate the median frequency of exchange, boxes indicate the upper and lower quartile values, and circles indicate outliers. Results are averaged over 500 simulations in each plot.

**Figure 6:**
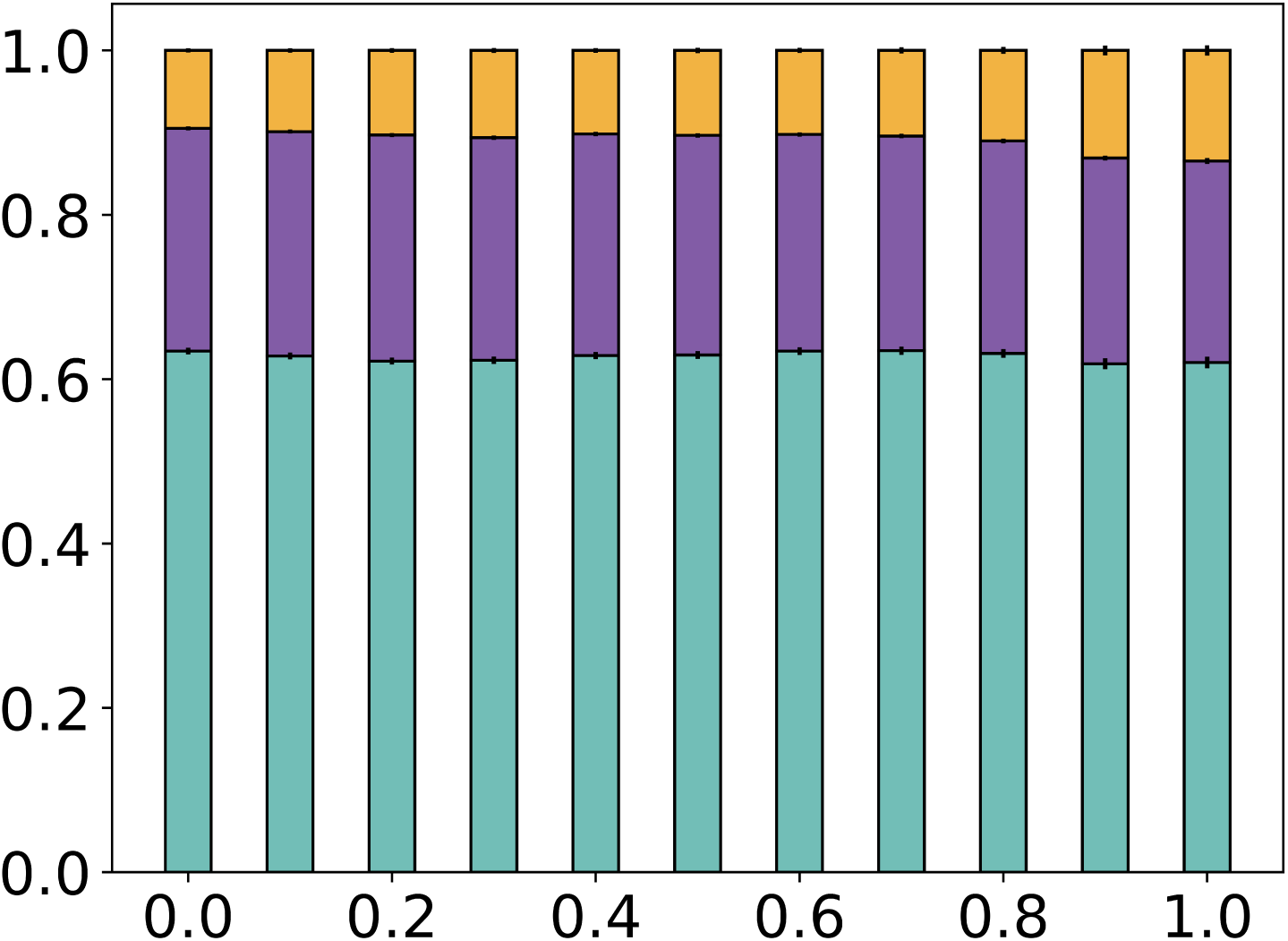
The confinement of migrating CNCCs to the r4-ba2 pathway as a function of the fraction of CNCC-free zones for which Colec12 is present in the original model representation of Colec12.

As before, migration occurs over a period of 24h, and the rules governing cell movement in the ABM are unchanged. In simulations, we characterise the exchange of a cell between streams as having occurred when a cell migrates from its original stream, crosses the mid-line of the CNCC-free zone adjacent to r3, and joins the neighbouring stream. For exchange to have occurred, this cell must then remain in this new stream until migration ends after 24h. We adopt this definition of cell exchange to prevent over-counting cell exchange when a CNCC fluctuates about the mid-line of CNCC-free zone adjacent to r3.

Model simulations indicate that the possibility of CNCC exchange between adjacent collectives depends primarily on the distance of separation between adjacent collectives (Figure 5C). In simulations, cells are exchanged between the streams adjacent to r2 and r4 when either Trail, Colec12, or Trail and Colec12 are present in the CNCC-free zone adjacent to r3. When only Colec12 is present, exchange occurs due to single-cell chemotaxis across the region adjacent to r3, when cells move into regions of Colec12 expression and extend filopodia into the adjacent stream where Colec12 is not expressed. However, in this context, we also find that both streams often develop diffuse edges which occasionally merge with one-another, disrupting migration (Supplementary Information S4). Conversely, when Trail is present in the CNCC-free zone (by itself or in addition to Colec12), the mechanism of CNCC exchange is different and leads to the formation of CNCC-bridges between the two streams without a large-scale disruption to migration. In this instance, the exchange of cells between the r4-ba2 and r2-ba1 pathways is observed when a cell loses directional information in its current stream due to a loss of cell-cell contact during migration (Figure 5B(i)). When this occurs, it is possible that filopodial protrusions extended by a cell proximal to the CNCC-free zone adjacent to r3 detect a cell in the adjacent stream (Figure 5B(ii)). If a cell in an adjacent stream is detected, then movement towards this cell is induced by the increase in cell-cell adhesion in regions of enhanced Trail expression within the CNCC-free zone adjacent to r3 (Figure 5B(iii)). Any other cells in this chain that also lie in the CNCC-free zone move towards the cell they are connected to, forming a chain that migrates across the CNCC-free zone (Supplementary Information S5).

If only Trail is expressed in the CNCC-free zone adjacent to r3, we find that CNCC bridges form consistently for stream separation distances of approximately 24*µ*m or less (Figure 5C), which corresponds to the approximate maximum radius of communication for a cell in the model (the maximum length of a filopodial protrusion). If both Trail and Colec12 are expressed in the CNCC-free zone adjacent to r3, the frequency of exchange is slightly lower than in the case of only Trail, as a smaller number of cells initially invade the region adjacent to r3. However, exchange occurs over a larger range of separation distances (approximately 36*µ*m), which corresponds to the increase in the length of filopodia extended by CNCCs exposed to Colec12 (Figure 5C). If only Colec12 is expressed in the CNCC-free zone adjacent to r3, exchange between streams occurs for a similar range of separations, but only due to single-cell migration down gradients in Colec12, and can lead to a disruption in migration when the diffuse edges of the two adjacent streams make contact.

Overall, simulations in this revised geometry suggest that CNCC exchange between adjacent streams can occur due to the hypothetical Trail and Colec12 mechanisms of action considered here. An increase in cell-cell adhesion in the presence of Trail may cause movement of cell chains across CNCC-free zones due to loss of contact guidance in a cell’s current stream together with connections with the adjacent stream. The morphological effects of Colec12 noted *in vitro* may increase the likelihood of contact with adjacent streams due to the formation of longer filopodial protrusions that span the region separating adjacent streams. Cells can also be exchanged between streams due to the effects of Colec12, though these exchanges represent single-cell migration across the CNCC-free zone, rather than the collective migration of chains observed *in vivo*. Here, we also note that Trail and Colec12 protein stripe assays analysed in prior studies of CNCC migration show cells stretching to lengths well beyond the maximum value prescribed in the ABM considered here [18]. However, CNCC lengths in these stripe assays were not precisely measured or quantitatively analysed, such that their values remain undetermined. If we were to reflect these observations in the ABM and increase the maximum radius of cell communication to larger values, then the width of the CNCC-free zone adjacent to r3 for which swapping occurs would likely increase to values comparable to the width of the CNCC-free zone adjacent to r3 *in vivo*. We highlight further study of the sizes of CNCCs exposed to Trail and Colec12 as a key experimental means of validating our proposed mechanism of CNCC exchange between adjacent streams.

## 3 Discussion

In this study, we used an ABM for CNCC migration to generate and test mechanisms for maintaining the coherence and spatial confinement of migrating CNCC streams. Our main objective was to study the effects of Trail and Colec12 on the confinement of CNCC streams to the stereotypical r4-ba2 pathway during migration. Prior *in vitro* studies have highlighted a change in the migratory dynamics and morphology of CNCCs cultured in the presence of Trail and Colec12 [18]. Furthermore, *in vivo* experiments have shown that loss-of-function of either Trail or Colec12 results in the increased diversion of chick CNCCs away from the stereotypical r4-ba2 migratory pathway into CNCC-free zones adjacent to r3 and r5. However, prior work in this context has not yet elucidated the mechanisms through which Trail and Colec12 confine CNCC streams to stereotypical migratory pathways *in vivo*. To this end, we conducted model simulations of CNCC responses to Trail and Colec12 mechanisms of action on CNCC behaviours derived from *in vitro* observations.

Model simulations predict that the presence of Trail in CNCC-free zones confines CNCC streams to stereotypical migratory pathways by enhancing cell-cell adhesion, resulting in the migration of CNCCs in the presence of Trail back towards areas of higher cell density in the r4-ba2 migratory pathway. Conversely, equivalent simulations of the presence of Colec12 in CNCC-free zones adjacent to the r4-ba2 pathway predict that the effects of Colec12 on the protrusive dynamics of CNCCs observed *in vitro* [18] have a negligible effect on the confinement of CNCCs to stereotypical migratory pathways in migration. We then hypothesised a mechanism of chemotaxis in CNCCs, motivated by observations that CNCCs exposed to Colec12 migrate back towards the neural tube in the chick, and found that this revised model increased the proportion of CNCCs confined to the r4-ba2 pathway throughout migration. As such, we predict that chemotaxis down gradients in Colec12 is the primary mechanism through which Colec12 restricts CNCC trajectories to stereotypical migratory pathways *in vivo* and that the morphological effects of Colec12 on CNCCs that have been observed *in vitro* are mechanisms that increase the likelihood of gradient detection in chemotaxis.

Model simulations also predict a synergy between Trail and Colec12, in that the confinement of CNCCs to stereotypical migratory pathways is maximised when both Trail and Colec12 are present in CNCC-free zones adjacent to the r4-ba2 pathway. In good agreement with prior studies [18], we also find that high levels of spatial confinement of CNCCs to the r4-ba2 pathway are achieved by expressing Trail and Colec12 in approximately the first third (≈ 400*µ*m) of the CNCC-free zones adjacent to r3 and r5. Collectively, model simulations suggest that both Trail and Colec12 are each factors that reduce the invasion of CNCCs into typically CNCC-free zones adjacent to r3/5. The Colec12 mechanism of action hypothesised here acts on individual cells to drive movement along external chemical gradients and re-connections with streams along the r4-ba2 pathway. Conversely, the Trail mechanism of action proposed here alters cell-cell interactions within collectives, and hence, primarily acts to keep CNCC streams as a whole within the region adjacent to r4. These two distinct mechanisms, therefore, maximise CNCC confinement in a complementary manner by driving both individual and collective CNCC migration towards the r4-ba2 pathway.

Previous experiments have highlighted the importance of Dan in collective chick CNCC migration [19]. *In vitro*, Dan has been found to modulate the speed at which CNCCs move. Moreover, *in vivo* experiments in the chick cranial neural crest have shown that Dan loss-of-function results in increased speed and directionality in CNCCs. Unlike Trail and Colec12, the expression profile of Dan extends parallel to the body axis all alongside r1-7 and, intriguingly, the concentration of Dan is reduced in regions through which leader CNCCs have previously migrated. In the ABM, we reflected these observations by representing Dan as a degradable factor expressed both along the stereotypical r4-ba2 migratory pathway, and in the regions void of CNCCs adjacent to r3 and r5. Simulations predict that while Trail and Colec12 prevent invasion into CNCC-free zones, they cannot prevent the frequent escape of cells at the leading edge of streams, resulting in the breakdown of directional guidance cues in streams and a subsequent separation of the stream. However, simulations in which Trail, Colec12, Dan, and VEGF are present in CNCC-free zones, and VEGF and Dan are present in regions adjacent to r3/4/5, suggest that Dan decreases the frequency of break-up of CNCC collectives by preferentially reducing the speed of cells at the leading edge of collectives to prevent their escape. In the model, Dan differentially modulates CNCC speeds through its degradation by CNCCs, in that cells at the leading edge of collectives are exposed to higher levels of Dan expression, and hence, have their speeds reduced by a greater amount than trailing cells that migrate through regions in which Dan has previously been degraded. Whilst previous models of CNCC migration have reproduced collective migration without considering the effects of Dan [12, 18, 21], successful migration in these models required VEGF degradation by CNCCs (and hence, the formation of VEGF gradients) to occur at sufficiently slow rates to prevent leader escape. The addition of Dan into the ABM considered here increased the range of feasible VEGF degradation rates for collective migration to occur. As such, we predict that both the expression of Dan along stereotypical migratory pathways and the degradation of Dan by CNCCs are mechanisms that confer robustness in collective invasion of the branchial arches. If Dan is expressed along stereotypical migratory pathways, and Trail/Colec12/Dan are expressed in adjacent CNCC-free zones, we find that CNCCs remain in streams confined to stereotypical migratory pathways that collectively invade the branchial arches for a wide range of VEGF degradation rates (and hence, maximum speed of movement of CNCCs at the leading edge).

A final prediction of the ABM is that Trail and Colec12 can drive a small number of cells to divert into CNCC-free zones and connect neighbouring streams through single cell-wide strands without disrupting the overall migratory pattern. The exchange of single cells between streams has previously been observed in the chick, where contact between CNCCs in adjacent streams is found to occur infrequently but consistently throughout migration [24]. Furthermore, in mice carrying null mutations of either npn2 or Sema3F, single-cell wide bridges have been observed to form between streams invading ba1 and ba2 [25]. Hitherto, this phenomenon has not been explained either with experiments or computational models. We predict that for adjacent collectives within approximately four CNCC radii (30*µ*m) of one-another (the approximate range of cell-cell interactions in regions of Colec12 expression), the effects of Trail and Colec12 on CNCC migratory behaviour facilitate contact between cells in adjacent streams. When a cell loses directional information in its current stream, it is possible that a cell in an adjacent stream is detected through the extension of filopodial protrusions. If this occurs, then cells in the presence of Trail begin to move towards the cell detected in an adjacent stream, leading to movement across a CNCC-free zone and into a neighbouring collective. We find that the exchange of CNCCs between adjacent streams increases as the distance between collectives is reduced, as contact between CNCCs in adjacent collectives occurs at a higher frequency. This effect is also observed when only Trail is present in CNCC-free zones, though adjacent streams must be closer to one-another for this to occur at the same frequency as when Colec12 is also expressed (due to the reduced range of interactions in CNCCs). We believe that this mechanism presents a possible explanation for the formation of CNCC bridges between adjacent collectives observed *in vivo* that is hitherto, without a mechanistic explanation.

Each of the biological hypotheses generated here merit further experiments to validate our predictions. Our hypothesis that Trail confines CNCC movement to stereotypical migratory pathways may be tested experimentally through studying the secretome of CNCCs cultured in the presence of Trail. If an upregulation of chemokines associated with cell clustering is observed in the secretome of CNCCs exposed to Trail, then it is likely that enhanced cell clustering is a primary mechanism through which Trail confines CNCCs to stereotypical migratory pathways *in vivo*. Furthermore, the exposure of CNCCs to Colec12 gradients in culture would provide a simple means of validating our hypothesis of CNCC chemotaxis down Colec12 gradients. Additionally, Trail and Colec12 stripe assays in which the width of Colec12/Trail-positive stripes separating adjacent CNCC streams are varied would allow for validation of the prediction that the distance at which adjacent CNCC streams are separated by regions of enhanced Trail and Colec12 expression is the major determinant in the formation of single-cell wide bridges between adjacent streams. It would also be desirable to measure the maximum observed cell length in these stripe assays, to determine if the putative mechanisms of CNCC exchange in the ABM are a sufficient mechanism for the exchange observed over longer distances *in vivo*. A final testable hypothesis generated by this model is that the degradation of Dan by CNCCs would result in a higher number of CNCC exchanges between adjacent streams in regions of the domain where exchanges have previously occured, due to a localised loss of speed modulation from Dan. This hypothesis could be easily tested in modified Trail and Colec12 protein stripe assays where Dan protein is also present.

It is important to acknowledge that these results were generated within a simplified model framework that makes multiple assumptions regarding CNCC migration and the geometry of the chick cranial neural crest. For example, we assumed the growth of the domain through which CNCCs migrate to be spatially uniform. However, recent *in vitro* co-culture assays have found mesoderm to proliferate at a higher rate in the presence of CNCCs [22]. It is, therefore, possible that the domain through which CNCCs migrate grows in a spatially non-uniform manner that depends on the local density of CNCCs. Future work will aim to reflect these findings and verify the robustness of our results to different spatial growth profiles of the domain through which CNCCs migrate. Moreover, the model uses a leader-follower representation of CNCC phenotypes, wherein cells are either a leader phenotype, with movement occurring solely via self-induced chemoattractant gradients, or a follower phenotype, where movement occurs solely through cell-cell guidance cues. This representation of CNCC phenotypes is motivated by observations of differential gene expression between chick CNCC leaders and followers [12]. However, it is likely that the molecular heterogeneities within a CNCC stream may vary in a continuous manner, rather than in the binary wavefront fashion considered here. In future studies, we will aim to refine the computational model to account for this (as in other modelling studies of neural crest migration [26]) to ensure the robustness of our findings to more sophisticated representations of CNCC phenotypes.

## 4 Experimental Procedures

### Computational model of cranial neural crest migration

We use a stochastic ABM written in Python3 to simulate the migration of chick CNCCs over a period of 24h. The model builds on prior ABMs of chick CNCC migration [12, 18, 19, 21] and is primarily parameterised as in [18]. We build on these models by integrating signalling affecting cell speed, direction, and cell-cell interactions into a single model framework. The full computational implementation of the model is described below.

#### 4.1 Domain growth

The domain through which CNCCs migrate is modelled as a two-dimensional rectangle of a fixed width (*L_y_* = 360*µ*m) and a length that increases in time. We model the growth of the domain through which CNCCs migrate to be spatially uniform, and logistic in time. Our implementation of domain growth is based on prior measurements in the chick cranial neural crest [12]. The length of the domain at time *t* during migration is given by

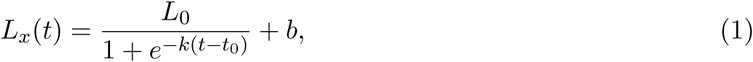

where *L*_0_ = 855.8*µ*m, *k* = 14.98h*^−^*^1^, *t*_0_ = 0.294h, and *b* = 294.3*µ*m. Parameter values were obtained by least squares fitting the experimental data to Equation (1).

#### 4.2 Chemical reaction-diffusion dynamics

We model the evolution of VEGF, Dan, Trail, and Colec12 concentration with four reaction-diffusion partial differential equations (PDEs) comprising terms for cell-induced degradation, production within the domain (assumed to be spatially uniform), dilution during domain growth, and diffusion. The concentration of chemical *j* (*j* ∈ {VEGF, Dan, Trail, Colec12}), *c_j_*(*x, y*), where (*x, y*) ∈ (0*, L_x_*(*t*)) × (0*, L_y_*), evolves according to

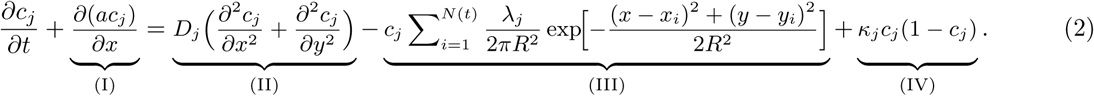

where term (I) corresponds to advection and dilution due to growth of the domain, term (II) represents diffusion, (III) is an internalisation term representing the local degradation of chemicals by cells (indexed by *i*), and term (IV) represents the spatially uniform production of the chemical within the domain. In solving Equation (2), we re-scale to a domain of unit length ((*x, y*) ∈ (0, 1) × (0*, L_y_*)) for numerical convenience. The re-scaled equation is given by

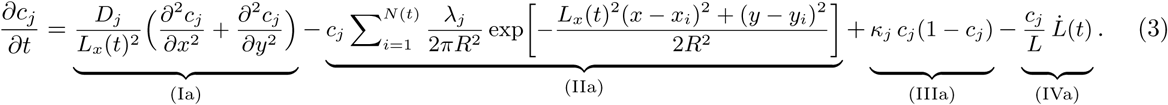

where term (Ia) represents diffusion, term (IIa) represents internalisation by cells, term (IIIa) represents spatially uniform production, and term (IVa) represents dilution due to domain growth. In simulations, we solve Equation (3) using a custom forward-Euler solver developed using Python3 (available at https://github.com/SWSJChCh/leaderFollower). To maintain numerical accuracy, we use time-steps smaller than those for cell movement (Δ*t*_solver_ = 0.1Δ*t*). We impose zero-flux boundary conditions on all chemicals at *x* = 0 and *x* = *L_x_*(*t*) and periodic boundary conditions on all chemicals at *y* = 0 and *y* = *L_y_*, to represent the presence of adjacent migratory pathways in the axial direction. The initial conditions are uniform concentrations of *c_j_*(*x, y*) = *c*_0_ = 1 in the regions where each chemical is expressed. Parameter names and values are given in Table 1, with some parameters set to zero to omit certain terms in Equation (3) for certain chemicals.

**Table 1:**
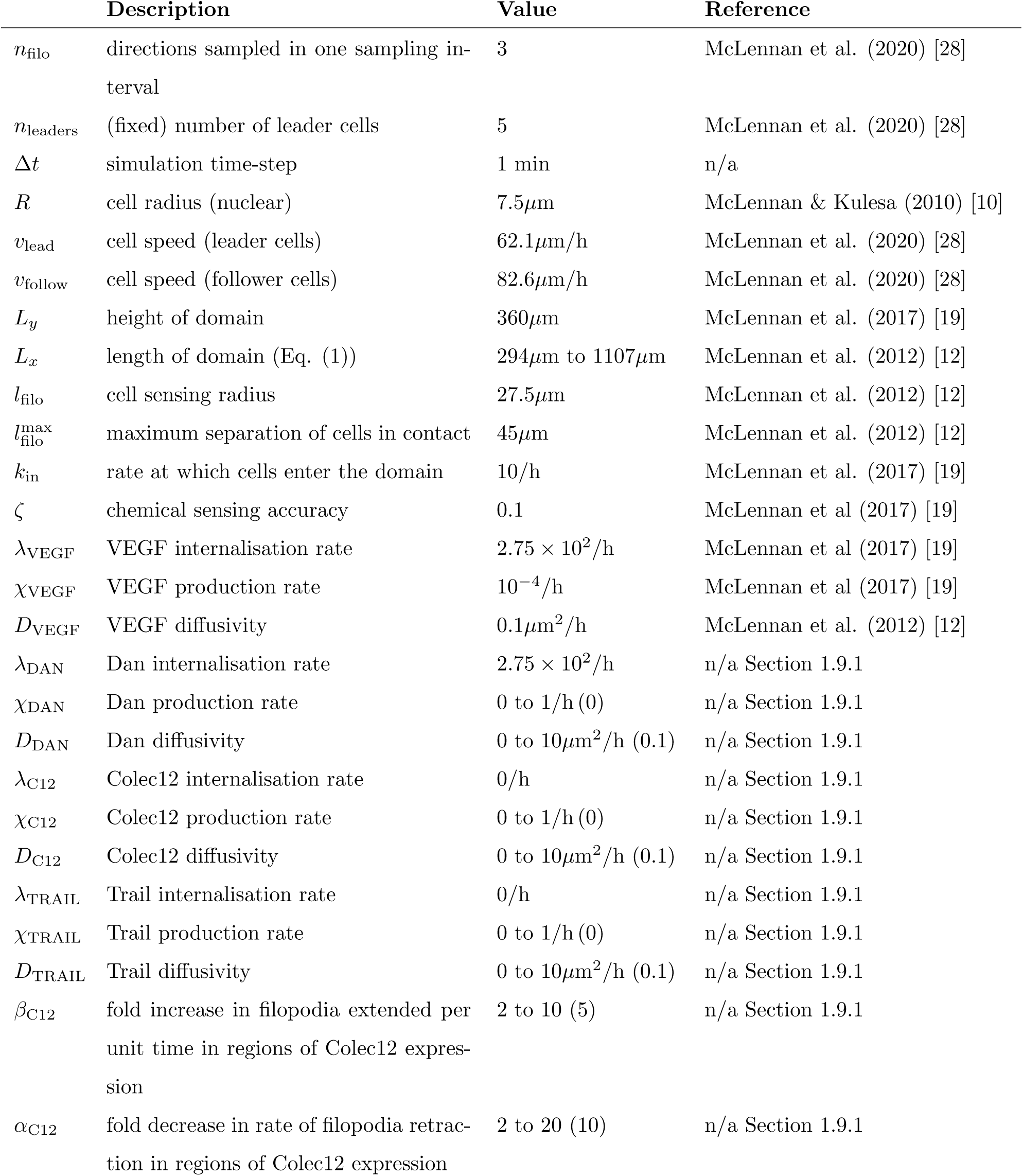
Default parameter values are given. When a range is given, model outputs are robust to a variation of the parameter in the stated range, and bracketed values were used as default. Chemical concentrations are given in relative units, such that for VEGF, Dan, Colec12, and Trail, *c*_0_ = 1.

|  | Description | Value | Reference |
| --- | --- | --- | --- |
| $n_{\text{filo}}$ | directions sampled in one sampling interval | 3 | McLennan et al. (2020) [28] |
| $n_{\text{leaders}}$ | (fixed) number of leader cells | 5 | McLennan et al. (2020) [28] |
| $\Delta t$ | simulation time-step | 1 min | n/a |
| $R$ | cell radius (nuclear) | $7.5\mu\text{m}$ | McLennan & Kulesa (2010) [10] |
| $v_{\text{lead}}$ | cell speed (leader cells) | $62.1\mu\text{m/h}$ | McLennan et al. (2020) [28] |
| $v_{\text{follow}}$ | cell speed (follower cells) | $82.6\mu\text{m/h}$ | McLennan et al. (2020) [28] |
| $L_y$ | height of domain | $360\mu\text{m}$ | McLennan et al. (2017) [19] |
| $L_x$ | length of domain (Eq. (1)) | $294\mu\text{m}$ to $1107\mu\text{m}$ | McLennan et al. (2012) [12] |
| $l_{\text{filo}}$ | cell sensing radius | $27.5\mu\text{m}$ | McLennan et al. (2012) [12] |
| $l_{\text{filo}}^{\text{max}}$ | maximum separation of cells in contact | $45\mu\text{m}$ | McLennan et al. (2012) [12] |
| $k_{\text{in}}$ | rate at which cells enter the domain | 10/h | McLennan et al. (2017) [19] |
| $\zeta$ | chemical sensing accuracy | 0.1 | McLennan et al (2017) [19] |
| $\lambda_{\text{VEGF}}$ | VEGF internalisation rate | $2.75 \times 10^2/\text{h}$ | McLennan et al (2017) [19] |
| $\chi_{\text{VEGF}}$ | VEGF production rate | $10^{-4}/\text{h}$ | McLennan et al (2017) [19] |
| $D_{\text{VEGF}}$ | VEGF diffusivity | $0.1\mu\text{m}^2/\text{h}$ | McLennan et al. (2012) [12] |
| $\lambda_{\text{DAN}}$ | Dan internalisation rate | $2.75 \times 10^2/\text{h}$ | n/a Section 1.9.1 |
| $\chi_{\text{DAN}}$ | Dan production rate | 0 to 1/h (0) | n/a Section 1.9.1 |
| $D_{\text{DAN}}$ | Dan diffusivity | 0 to $10\mu\text{m}^2/\text{h}$ (0.1) | n/a Section 1.9.1 |
| $\lambda_{\text{C12}}$ | Colec12 internalisation rate | 0/h | n/a Section 1.9.1 |
| $\chi_{\text{C12}}$ | Colec12 production rate | 0 to 1/h (0) | n/a Section 1.9.1 |
| $D_{\text{C12}}$ | Colec12 diffusivity | 0 to $10\mu\text{m}^2/\text{h}$ (0.1) | n/a Section 1.9.1 |
| $\lambda_{\text{TRAIL}}$ | Trail internalisation rate | 0/h | n/a Section 1.9.1 |
| $\chi_{\text{TRAIL}}$ | Trail production rate | 0 to 1/h (0) | n/a Section 1.9.1 |
| $D_{\text{TRAIL}}$ | Trail diffusivity | 0 to $10\mu\text{m}^2/\text{h}$ (0.1) | n/a Section 1.9.1 |
| $\beta_{\text{C12}}$ | fold increase in filopodia extended per unit time in regions of Colec12 expression | 2 to 10 (5) | n/a Section 1.9.1 |
| $\alpha_{\text{C12}}$ | fold decrease in rate of filopodia retraction in regions of Colec12 expression | 2 to 20 (10) | n/a Section 1.9.1 |

#### 4.3 Sensing accuracy

Our implementation of the sensing accuracy of chemotactic gradients by cells is based on prior work by Berg and Purcell [27] and is as described in prior models of chick CNCC migration [19]. In the ABM, a filopodial protrusion senses a chemical gradient when

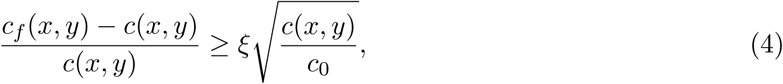

where *c_f_* (*x, y*) is the average chemical concentration integrated along the filopodium, *c*(*x, y*) is the chemical concentration at a cell’s current position, *c*_0_ is the initial uniform concentration of the chemical within the domain, and *ξ* is a dimensionless parameter that determines the accuracy of chemotactic gradient detection by cells. Parameter names and values are given in Table 1.

#### 4.4 Cell movement in the presence of Dan

We assume that the speed of migrating CNCCs is modulated in the presence of Dan protein according to

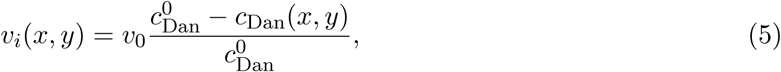

where *v_i_*(*x, y*) is the speed of movement of CNCC *i* at a position (*x, y*), *v*_0_ is the maximum speed of movement for CNCC *i* (dependent on its phenotype), 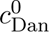 is the initial (uniform) concentration of Dan in the domain, and *c*_Dan_(*x, y*) is the concentration of Dan at a position (*x, y*) in the domain. Equation (5) means that CNCC movement speed decreases linearly with the local concentration of Dan, and that movement ceases completely in regions where Dan is of an initial maximum concentration of 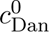. A similar representation of CNCC speed modulation in regions of Dan expression was considered in a prior model of CNCC migration [19], though in this model, Dan was not degraded by CNCCs, and hence was of a spatially uniform concentration in the domain that varied as a function of time.

#### 4.5 Cell movement in the presence of Trail

We represent the effects of Trail protein on CNCCs *in vitro* through multiple modifications to the behaviour of CNCCs. When a CNCC encounters Trail, its speed is reduced by 25%, in line with *in vitro* observations [18]. To represent increased cell-cell adhesion in the presence of Trail, we modify the primary mechanism of movement for CNCCs in chains. When not in the presence of Trail, follower cells in our model move by forming chains of cells with a leader at the front (Section S1.8). All cells in a given chain move in the same direction as the leader cell at the front of the chain. In CNCCs that encounter Trail, we modify this mechanism of movement, such that follower cells instead move towards the cell directly in front of them in their respective chain, meaning that movement in the presence of Trail is directed towards areas of higher cell density, on average.

#### 4.6 Cell movement in the presence of Colec12

We represent the effects of Colec12 protein on CNCCs *in vitro* through multiple modifications to the behavior of CNCCs. When a CNCC encounters Colec12, the length of filopodial protrusions it extends is increased by 50%, in line with *in vitro* observations [18]. Furthermore, we represent the increased branching of filopodia by the extension of a higher number of filopodia in a given time interval (Section S1.9.1). To represent the slower rate of filopodia retraction in CNCCs exposed to Colec12 protein, we also increase the time interval between the retraction and extension of new filopodia for CNCCs, such that a given filopodium remains extended for a longer time interval (Section S1.9.1). In simulations where the effects of Colec12 protein on CNCCs were limited to the morphological changes observed *in vitro*, the presence of Colec12 in CNCC-free zones adjacent to r3/5 was insufficient to confine CNCCs to the r4-ba2 pathway (Section S2). As such, we also added a mechanism of chemotaxis, in which CNCCs migrate down gradients in Colec12 protein and back towards the r4-ba2 pathway where its concentration is lower. The detection of gradients in Colec12 by CNCCs is a modification of Equation (4) in S1.3, such that a gradient in Colec12 is detected if

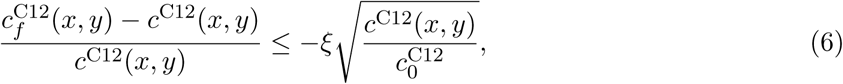

where *c*^C12^(*x, y*) is the average concentration of Colec12 integrated along the filopodium, *c*^C12^(*x, y*) is the concentration of Colec12 at a cell’s current position, *c*^C12^ is the initial uniform concentration of Colec12 within the domain, and *ξ* is a dimensionless parameter that determines the accuracy of Colec12 gradient detection by cells.

#### 4.7 Phenotype switching

Our implementation of phenotype switching is similar to that of a prior model for CNCC migration [18]. *In vivo*, VEGF signaling has been found to induce the expression of genes associated with leader cell behaviour in the chick cranial neural crest [21]. To reflect these observations in the ABM, we approximated this mechanism by fixing the number of leader cells in each simulation (Section S1.9), and updating CNCC phenotypes after movement according to their position in the domain. If, after movement, a follower cell lies a distance *ɛ* ahead of a leader cell, their phenotypes are switched, such that the follower cell adopts a leader phenotype and vice-versa. If a follower cell lies a distance *ɛ* ahead of multiple leader cells after movement, it switches phenotypes with the leader cell it is closest to. This mechanism ensures that CNCCs with a leader phenotype are those which lie in the regions of highest VEGF concentration along the r4-ba2 pathway, on average.

#### 4.8 Model pseudocode

##### 4.8.1 Main ABM loop

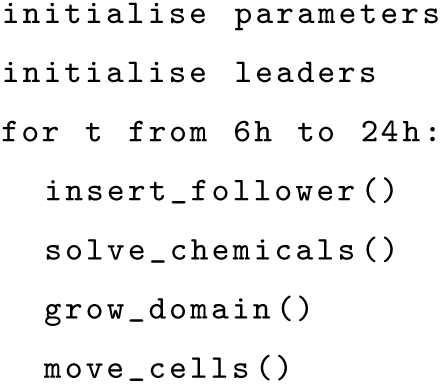

##### 4.8.2 Main loop functions

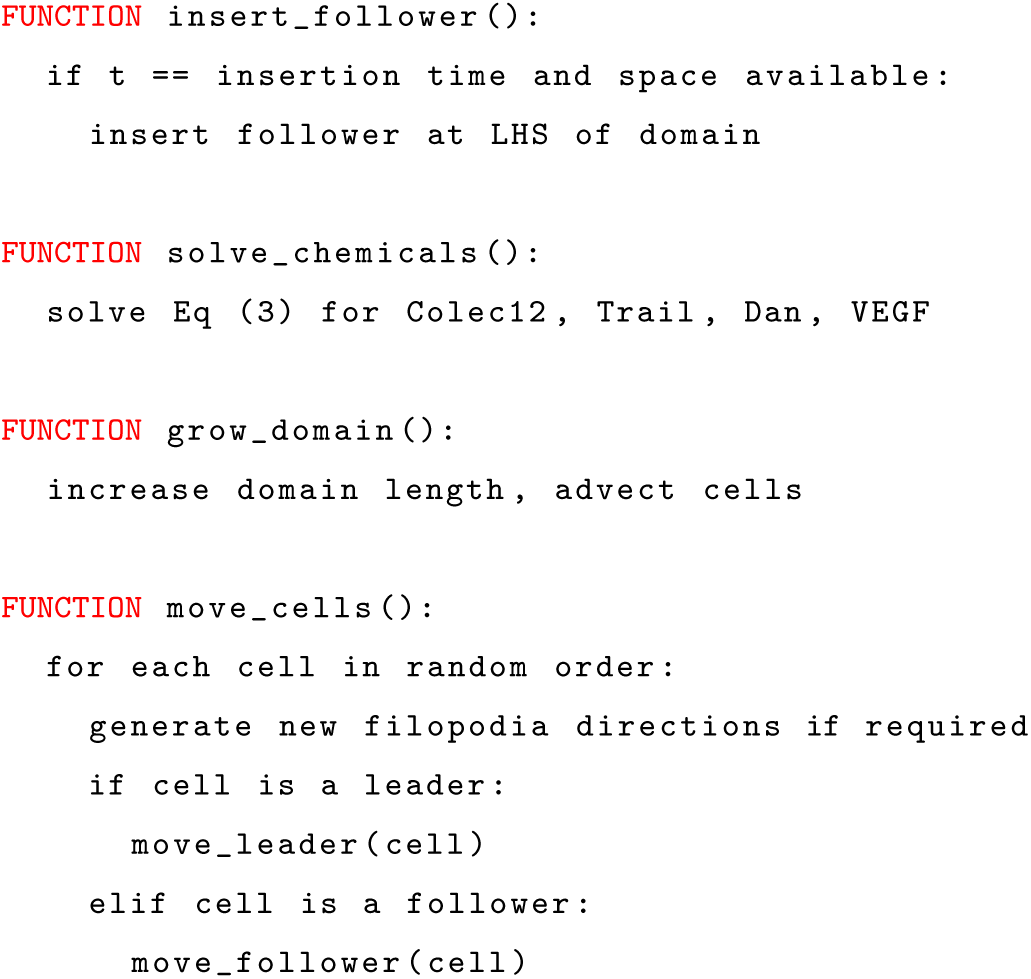

##### 4.8.3 Helper functions

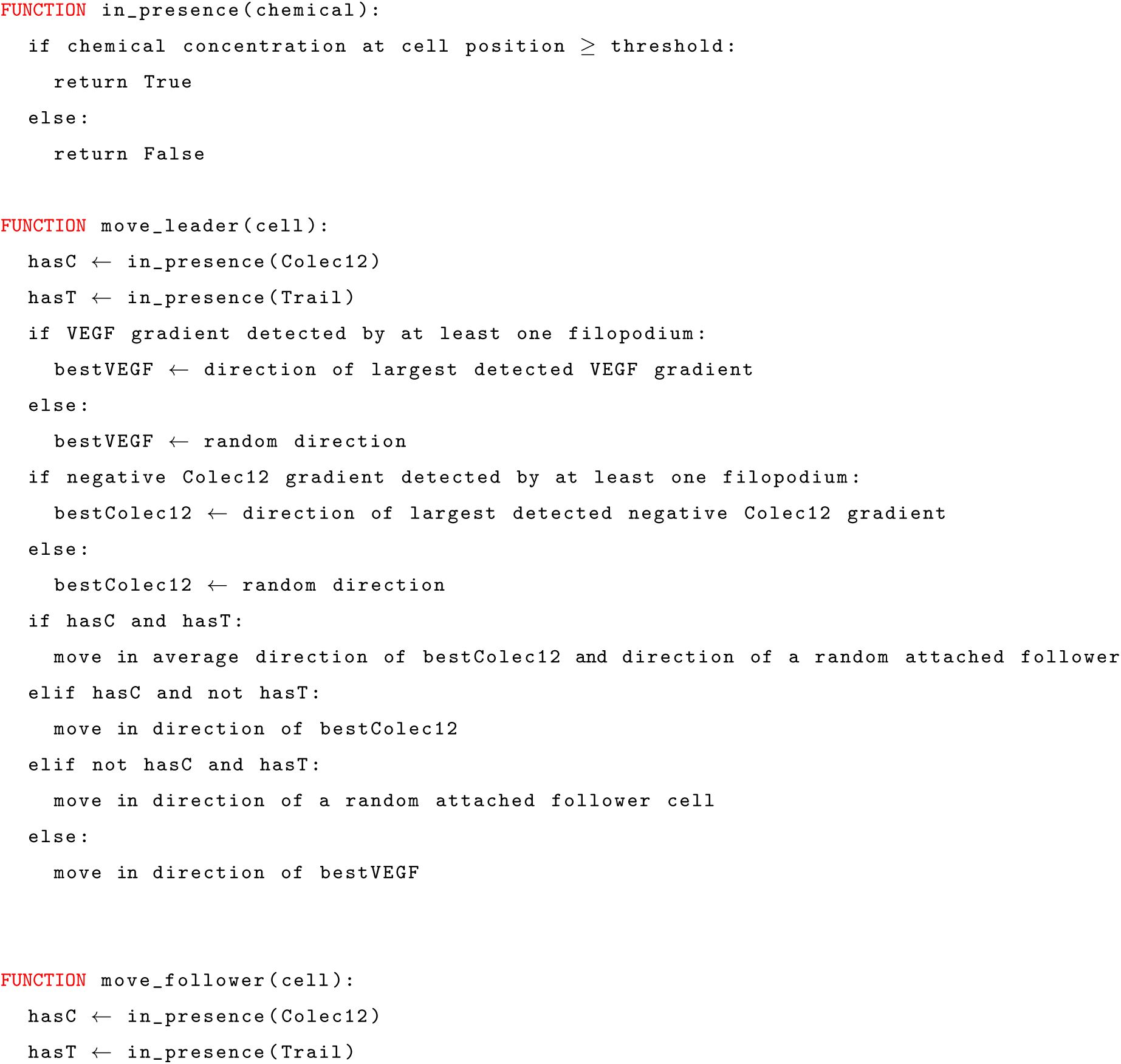
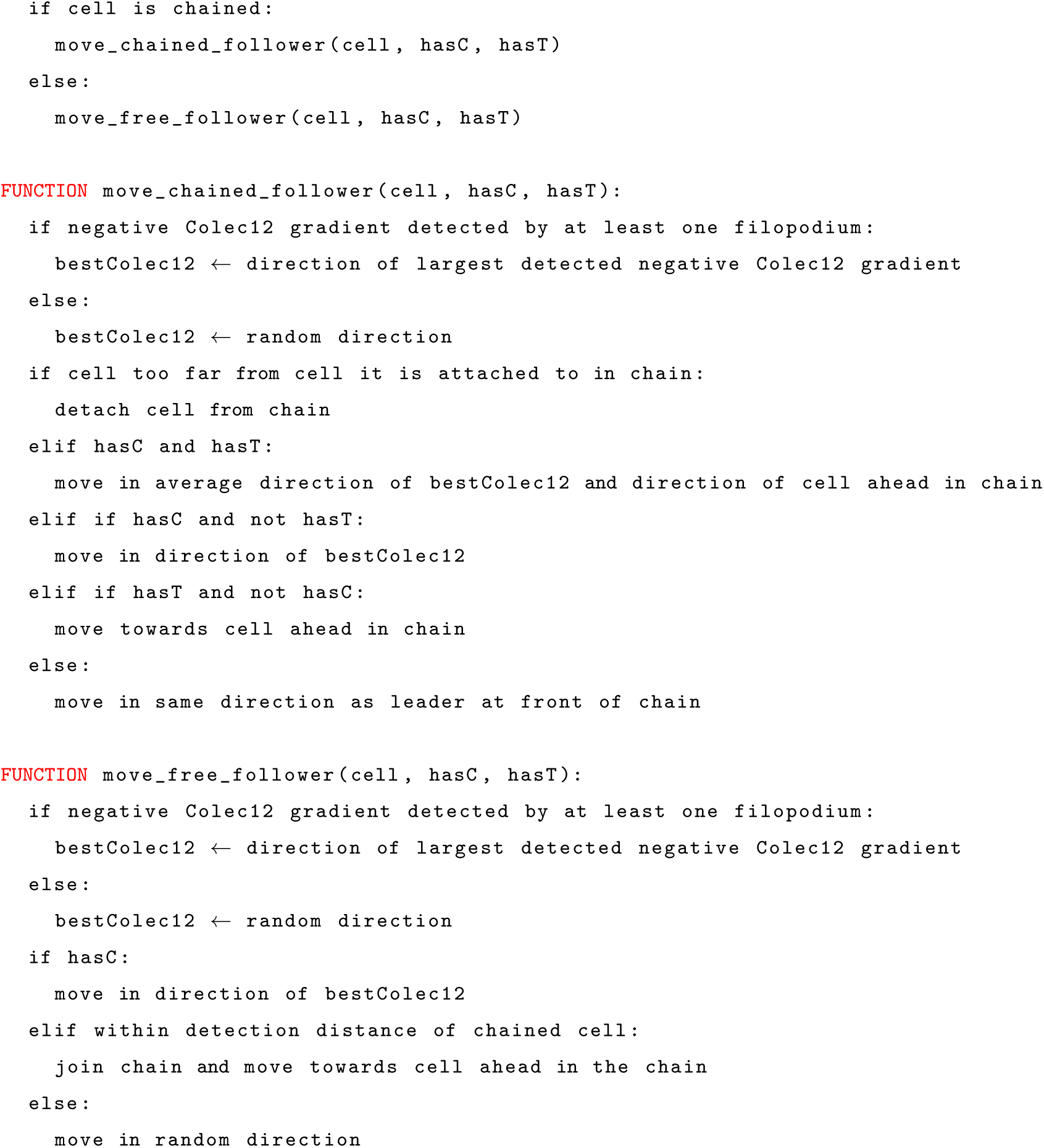

#### 4.9 Model parameterisation

We detail the parameters used in ABM simulations (Table 1), and briefly discuss the values of select parameters.

##### 4.9.1 Notes on model parameterisation

**Diffusivity of Trail, Colec12, and Dan**

The diffusivity of Trail, Colec12, and Dan have not been quantified experimentally. In the absence of experimental parameterisation, we assume that the diffusivity of Colec12, Trail, and Dan are the same as that of VEGF. In general, we find that VEGF, Trail, Colec12, and Dan must all be of a diffusivity

##### 4.9.2 Internalisation rate of Trail, Colec12, and Dan

The degradation of Trail and Colec12 by CNCCs has not been observed *in vivo*. As such, CNCCs in the model do not degrade Trail or Colec12 as they migrate. However, in the chick neural crest, Dan has been observed to exhibit a dynamic expression profile during CNCC migration, in that Dan concentration is lower in regions through which CNCCs have previously migrated, when compared with regions through which migration has not occured [19]. As such, we hypothesise that CNCCs degrade Dan during migration. In the absence of experimental studies that allow us to parameterise this rate of degradation, we assume Dan to be degraded at the same rate as VEGF (2.75 × 10^2^*/*h).

##### 4.9.3 Production rate of Trail, Colec12, and Dan

There is no *in vivo* evidence to suggest that Dan, Trail, and Colec12 are produced in the paraxial mesoderm during CNCC migration. As such, our default production rates of Dan, Trail, and Colec12 are *χ*_Dan_ = *χ*_TRAIL_ = *χ*_C12_ = 0*/*h. However, our findings are robust for all values of *χ*_TRAIL_ and *χ*_C12_ with *χ*_Dan_ ≲ 1*/*h.

##### 4.9.4 Fold increase in the number of filopodia extended when CNCCs encounter Colec12

*In vitro*, CNCCs in the presence of Colec12 exhibit increased branching in the filopodia they extend when compared with control media [18]. We represent increased branching in filopodia by the extension of a higher number of filopodia in regions of enhanced Colec12 expression. There is no way to directly link the increased branching observed *in vitro* to the fold increase in filopodia extended in our model. As a default value, the fold increase in filopodia extended by CNCCs that encounter Colec12 protein expression is *β*_C12_ = 2, but our findings are robust to large variations in this parameter (*β*_C12_ = 2 to 10).

##### 4.9.5 Fold decrease in the rate of filopodia retraction when CNCCs encounter Colec12

*In vitro*, CNCCs in the presence of Colec12 retract filopodia at a slower rate than in control media [18]. However, the fold decrease in the rate at which filopodia are retracted in regions of Colec12 expression has not been quantified experimentally. We chose a default value of *α*_C12_ = 10, but our findings are robust to large variations in this parameter (*α*_C12_ = 2 to 20).

#### 4.10 Original model representation of Colec12

The original model representation of the interaction between CNCCs and Colec12 protein was based on *in vitro* observations that CNCCs cultured in the presence of Colec12 extend longer, more branched filopodia that our retracted at a slower rate [18]. We represented these observations in the ABM by having CNCCs that encounter Colec12 extend twice the number of filopodia, each of 1.5 times the length at baseline. Furthermore, CNCCs retracted filopodia at a rate 10 times slower than at baseline in the presence of Colec12. Model simulations in which the proportion of CNCC-free zones adjacent to r3/5 for which Colec12 was present suggested that these effects of Colec12 were insufficient to confine CNCC migration to the r4-ba2 pathway. For all distances of Colec12 expression in the *x* direction, we found that the endpoint of around 35% of CNCC trajectories lay outside of the r4-ba2 pathway after 24h. The results in this figure are averaged over 500 simulations.

## Dedication

This paper was submitted for publication just weeks before the untimely passing of Paul M. Kulesa.

Paul gained his doctorate in mathematical biology under the supervision of Professor J.D. Murray. As a theoretician, Paul felt frustrated that model predictions were rarely tested so he trained as an experimentalist under the guidance of Professor Scott Fraser. This gave him the powerful set of combined skills in being able to determine which biological questions were amenable to mathematical modelling and which were not. Thus began a fruitful collaboration between us and the Kulesa laboratory that spanned over a decade. During this time, we combined theory with experiment to unearth a number of key findings in neural crest biology. Paul was an excellent scientist, always full of new ideas and very passionate about combining mathematics with biology to lead to insights that could not be achieved by either discipline alone. He was also very warm, kind and supportive, with a cheeky sense of humour that always made us laugh. He leaves behind his wife, Jennifer Kasemeier-Kulesa, who is also a Senior Researcher in his lab, and five children. Paul will be sorely missed by all who knew him, but he leaves behind a rich legacy of scientific work that will live long into the future.

## Data and code availability statement

Code used in the generation of the data in this study can be found at https://github.com/SWSJChCh/ leaderFollower.

## Authors’ contributions

**Samuel Johnson:** Data curation (lead); formal analysis (lead); methodology (lead); software (lead); writing–original draft (lead); writing–review and editing (supporting) **Paul M. Kulesa**: Conceptualisation (equal); investigation (equal); software (supporting); supervision (supporting); writing–original draft (supporting); writing–review and editing (lead). **Ruth E. Baker**: Conceptualisation (equal); investigation (equal); software (supporting); supervision (lead); writing–original draft (supporting); writing–review and editing (lead). **Philip K. Maini:** Conceptualisation (equal); investigation (equal); software (supporting); supervision (equal); writing–original draft (supporting); writing–review and editing (lead).

## Acknowledgements

SJ receives support from the Biotechnology and Biological Sciences Research Council (BBSRC) (grant number BB/T008784/1). R.E.B. is supported by a grant from the Simons Foundation (MP-SIP-00001828). For the purpose of open access, the author has applied a CC BY public copyright licence to any author accepted manuscript arising from this submission.

